# Transcription Factor Regulation of RNA polymerase’s Torsional Capacity

**DOI:** 10.1101/306258

**Authors:** Jie Ma, Chuang Tan, Xiang Gao, Robert M. Fulbright, Jeffrey W. Roberts, Michelle D. Wang

## Abstract

During transcription, RNA polymerase (RNAP) supercoils DNA as it forsward-translocates. Accumulation of this torsional stress in DNA can become a roadblock for an elongating RNAP and thus should be subject to regulation during transcription. Here, we investigate whether, and how, a transcription factor may regulate the torque generation capacity of RNAP and torque-induced RNAP stalling. Using a real-time assay based on an angular optical trap, we found that under a resisting torque, RNAP was highly prone to extensive backtracking. However, the presence of GreB, a transcription factor that facilitates the cleavage of the 3’ end of the extruded RNA transcript, greatly suppressed backtracking and remarkably increased the torque that RNAP was able to generate by 65%, from 11.2 to 18.5 pN·nm. Analysis of the real-time trajectories of RNAP position at a stall revealed the kinetic parameters of backtracking and GreB rescue. These results demonstrate that backtracking is the primary mechanism that limits transcription against DNA supercoiling and the transcription factor GreB effectively enhances the torsional capacity of RNAP. These findings broaden the potential impact of transcription factors on RNAP functionality.

## INTRODUCTION

RNA polymerase (RNAP) is a powerful biological motor, capable of generating both forces and torques. Although the force generation capacity has been well characterized for both *E. coil* RNAP (1) and Pol II (2), much less is known about RNAP’s capacity to generate torque (3) (Supporting Information Text). During transcription, RNAP over-twists DNA in the front (downstream) and under-twists DNA behind (upstream), as described by the twin-supercoiled domain model (4). Increasing evidence is emerging that transcription induces genome-wide supercoiling that transmits thousands of base pairs from transcription start sites (5–7) and the accumulation of torsional stress may hinder further translocation of RNAP if not relaxed in a timely fashion via DNA rotation or through the action of topoisomerases. Such torsion may also up– or down-regulate transcription by creating highly supercoiled DNA, encouraging the formation of non-B DNA (8, 9), recruiting regulatory proteins (10–12), and/or removing roadblock proteins (13).

Although RNAP generated torsion is widely recognized as an essential component of fundamental cellular processes and has been shown to have significant impact on gene expression, the torsional properties of RNAP have yet to be fully understood. As a torsional motor, RNAP may be characterized by its torque generation capacity, and this valuable parameter was long sought after (4, 11, 12, 14–16). Historically, the most prevalent approach has been biochemical studies using two-dimensional gels which have provided a wealth of information on the degree of DNA supercoiling generated by transcription. Recent torque measurements during real-time transcription have allowed a more complete understanding of the torsional mechanics. In particular, studies using the angular optical trap (AOT) showed that excessive torque accumulation during transcription stalls an elongating *E. coli* RNAP and the measured stall torque of RNAP is ^∼^11 pN·nm (17). This value represents the intrinsic capacity of RNAP for torque generation and establishes a baseline for a physiologically relevant torque scale.

Intriguingly, under (-) DNA supercoiling behind a transcribing RNAP, the mean stall torque value, 11 pN·nm, coincides with the torque required for DNA melting, where DNA undergoes a phase transition from B-DNA to melted, denatured DNA (18–20), and this transition plays critical roles in gene regulation, such as facilitating transcription initiation (21–23) and binding of transcription factors (11, 12). Conversely, (+) DNA supercoiling ahead of the RNAP may dissociate protein road blocks (13) and thus facilitate the passage of an elongating RNAP. Thus, perturbations to the torque generation capacity should lead to sensitive tuning of the level of gene expression. Although under (-) supercoiling, the torque RNAP can generate is limited to ^∼^ 11 pN·nm due to the DNA melting transition, under (+) supercoiling DNA is able to sustain a much greater torque without undergoing any structural denaturation (24–27). However, it is unclear whether and how RNAP may further increase its torque generation capacity under (+) supercoiling.

*In vivo*, RNAP functions closely with many transcription factors which regulate RNAP activities (28–30), therefore it is possible that these transcription factors might also regulate the torsional properties of RNAP. Our previous studies suggested that torque-induced stalling might be due to backtracking (17), during which RNAP reversely translocates along DNA with the catalytic site disengaged from the 3’-end of the RNA, inactivating transcription (31, 32). A universal class of transcription elongation factors, including TFIIS in eukaryotic cells (33–34) and GreB in prokaryotic cells (35, 37), rescues backtracked complexes and promotes transcription through obstructive regions of DNA such as regulatory transcription pause sites. These proteins act either as auxiliary factors or in the case of RNA Polymerase III, as intrinsic parts of the core RNAP (38). They stimulate the intrinsic cleavage activities of RNAP, leading to the removal of the 3’-end of the RNA. The newly generated RNA 3’-end then becomes aligned with the catalytic site, reactivating transcription. Thus here we have systematically investigated whether and how *E. coli* GreB interaction with RNAP allows RNAP to more efficiently transcribe through regions of DNA under torsional stress.

## RESULTS

### Single Molecule Assay for Transcription Stalling under Torque

To investigate RNAP’s stall torque regulation, we must first fully understand the nature of the torque-induced stalling. We thus employed a single molecule AOT assay (17, 26, 39) to closely examine transcription stalling in real time under (+) DNA supercoiling in the presence or absence of the transcription factor GreB.

As shown in Fig. 1A, an RNAP was torsionally anchored to the surface of a microscope coverslip, with the downstream end of its transcribing DNA torsionally anchored to the bottom of a nanofabricated quartz cylinder held in an AOT. Subsequently, RNAP elongation was directly monitored as torque accumulated on downstream DNA (Fig. 1B and 1C). Under this configuration, RNAP forward translocation introduced (+) supercoiling into the DNA, buckling the DNA to form a plectoneme and inducing a rapid shortening of DNA extension. As torque increased, RNAP was ultimately stalled, which we define here as < 1 bp/s forward translocation for approximately 60 s. This velocity threshold is significantly below the pause-free elongation velocity under zero torque (^∼^20 bp/s) (1, 40) while being well above the drift velocity of the instrument (^∼^ 0.06 bp/s) (Fig. S1). In addition, the force applied here (< 1.5 pN) was much smaller than the force required to stall the RNAP (about 25 pN) (1, 41) and thus stalling was primarily a result of the resisting torque.

**Figure 1.**
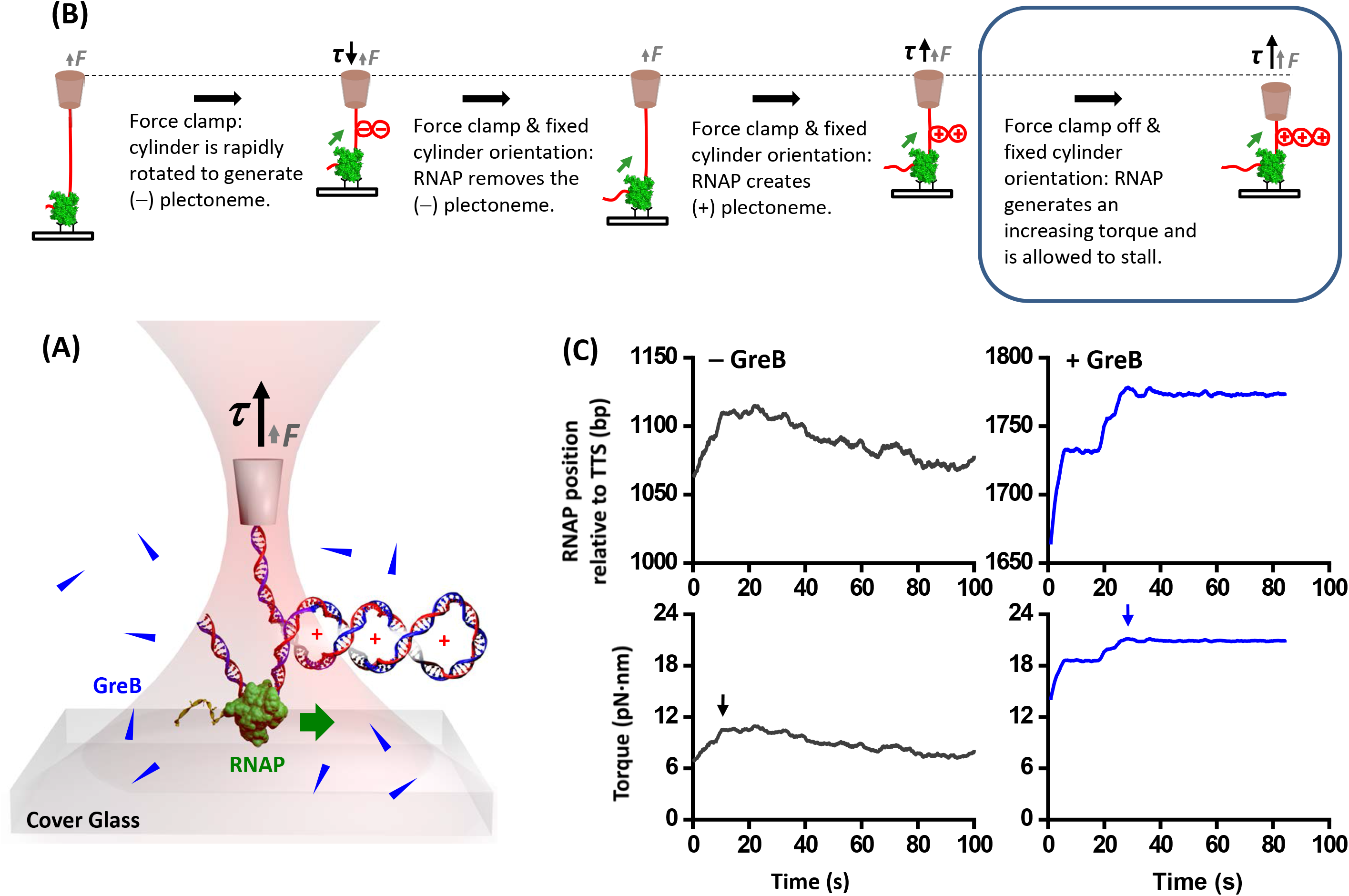
Measuring RNAP stall torque in the presence of GreB. (**A**) Experimental configuration of transcription stalling under torsion using an angular optical trap. RNAP was torsionally anchored to the surface of a microscope coverslip. A quartz cylinder was attached to the downstream end of the DNA and was held in the angular optical trap. RNAP transcription induced (+) supercoiling in the downstream DNA. (**B**) Experimental steps of the stall torque measurements. Under a constant force, the DNA was mechanically under-wound to generate a (-) supercoiling induced plectoneme. Subsequent RNAP forward translocation removed the (-) supercoiling, introduced (+) supercoiling, and buckled the DNA to form a (+) plectoneme. At this point, force feedback was turned off so that both the coverslip and trap center positions were held fixed. Further transcription led to an increase in the force and a corresponding increase in the buckling torque, eventually resulting in transcription stalling. (**C**) Representative traces of RNAP stall torque measurements in the absence or presence of GreB during the last step of stalling experiments. RNAP template position was defined as the RNAP’s physical position relative to the transcription start site (TTS). Arrows indicate the beginning of stalling.

### GreB Limits RNAP Backtracking after a Torque-Induced Stall

To characterize RNAP behavior at a stall, we examined RNAP movement during stalling by characterizing its maximum backtracking distance which is defined as the maximum reverse distance during the 60 s of stalling. In the absence of GreB (Fig. 1C), RNAP became significantly backtracked upon stalling. The mean maximum backtracking distance was −40.4 ± 4.8 bp (mean ± SEM) with 70% backtracked over 20 bps and some even exceeding 100 bp. The extent of backtracking is significantly greater than force-induced backtracking reported earlier for *E. coli* RNAP (1, 42–44). These results demonstrate that a resisting torsional stress can be highly effective in inducing RNAP backtracking.

In contrast, the presence of 1 μM GreB greatly reduced the backtracking distance. Approximately 94% of traces showed a maximum backtracking distance of −9.7 ± 1.1 bp (mean ± SEM) (Fig. 1C; Fig. 2B), which is a four-fold reduction from that seen with RNAP alone. The remaining 6% of traces showed a much greater backtracking distance of around 40 bp, comparable to those of RNAP alone. We interpret these traces as corresponding to RNAP molecules that were unable to interact with GreB. Indeed, a recent single molecule study of GreB by fluorescence visualization also found that a minority fraction of RNAP molecules was incapable of binding to GreB (45). We thus excluded this fraction from further analysis.

**Figure 2.**
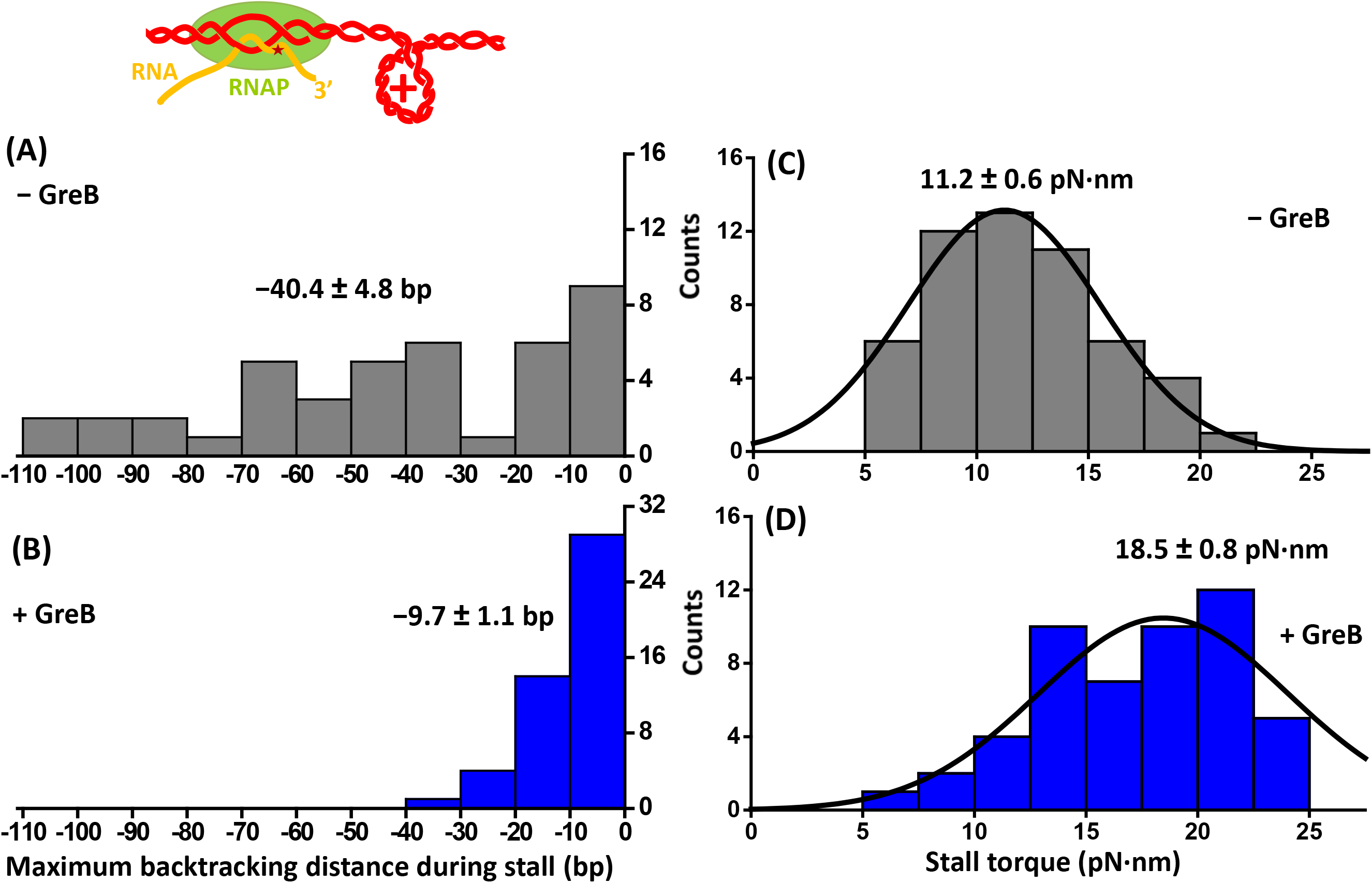
RNAP backtracking distance and stall torque in the absence and presence of GreB. (**A**) The distributions of the maximum backtracking distance during stalling in the absence and presence of GreB. For each distribution, the mean value and its SEM are indicated. (**B**) The distributions of the stall torque in the absence and presence of GreB. Each distribution was fit with a Gaussian function. For each distribution, the mean value of the Gaussian fit and its SEM were indicated.

These results demonstrate that, even when RNAP is under substantial torsion, GreB can still access the secondary channel of the RNAP, interact with the RNAP, and effectively limit backtracking. Upon RNAP backtracking, a bound GreB may stimulate the intrinsic transcript cleavage activity of RNAP, resulting in the 3’-OH of the newly cleaved RNA being aligned with the catalytic center of the enzyme. Subsequent resumption of transcription provides a renewed opportunity for RNAP to work against a greater torque.

### GreB Greatly Enhances Stall Torque Capacity

To examine the effect of GreB on the torque generation capacity of RNAP, we directly measured the stall torque of RNAP, both in the absence and presence of GreB. For each trace, the torque at which RNAP stalled was determined (Fig. 1C) and pooling data from multiple traces yielded a stall torque histogram. In the absence of GreB, the stall torque distribution was fit by a single Gaussian function, yielding a mean torque of 11.2 ± 0.6 pN·nm (mean ± SEM) (Fig. 2C), consistent with our previous measurements (17). In the presence of GreB, the stall torque distribution shifts to greater values with a mean torque of 18.5 ± 0.8 pN·nm, 65% greater than the stall torque generated by RNAP alone.

These data clearly demonstrate that GreB can play an important role in up-regulating RNAP’s torsional capacity during transcription. Although the torque enhancement is attributable to the presence of GreB, torque generation capacity of RNAP stems from the RNAP itself as GreB is known to merely serve as a catalyst in the conversion of a backtracked complex to an active elongation complex.

### Kinetics of RNAP Backtracking and RNA Cleavage

To better understand the stalling behavior in the presence of GreB, we must determine the kinetic rates of backtracking and GreB rescue during a stall. The trajectories of the RNAP position provide critical information relating to these kinetic parameters. The process of torque-induced stalling represents a competition between backtracking and rescue. We thus modeled this process as reversible first-order transitions between two-states (backtracking and elongating). The lifetimes of the two states and the velocities of RNAP in the two states determine the time course of RNAP’s position. Using rigorous techniques based on statistical mechanics and Eigen-vector analysis (Supplementary Information Text), we obtained analytical expressions for both the mean RNAP position and the variance of the RNAP position in the presence of GreB at a stall as a function of time 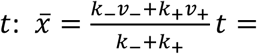 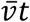 (Eq. S17) and 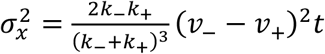 (Eq. S19) respectively. These quantities are expressed in terms of four kinetic parameters: the backtracking velocity *v_-_*, the active elongation velocity *v_+_*, the GreB rescue rate *k_+_*, and the rate to enter backtracking *k_-_*. Thus *k_-_* and *k_+_* may be determined as long as 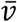 (mean RNAP velocity), *v_-_*, and *v_+_* can be measured.

We determined the backtracking velocity *v_-_* from the real-time motion of RNAP during a stall in the absence of GreB. These data also provide a unique window of opportunity to dissect the backtracked states. Fig. 3A (top) shows representative traces with a torque ^∼^ 18 pN·nm upon stalling, close to the mean stall torque with GreB; RNAP’s reverse motion due to backtracking was readily detectable. Because GreB was absent, RNAP underwent extensive backtracking without any rescue. The backtracking velocity increased with torque (Fig. 3B, top). We model the backtracking process as a one-dimensional Brownian diffusion process (40, 46, 47), with an effective hopping rate *k*_0_ between backtracking states in the absence of torque and the net motion biased by torque (Supplementary Information Text). Analysis of the RNAP velocity 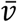 dependence on torque yielded *k*_0_ = 0.26 s^-1^. Thus we were able to make a rather direct measurement of the backtracking hopping rate. To our knowledge, this rate has not been determined for *E. coli* RNAP. Our value is comparable to those estimated for Pol II using pause duration analysis (47). Our analysis of *E. coli* RNAP yielded a backtracking velocity of *v_-_* = −0.9 bp/s at 18.5 pN·nm (the measured mean stall torque with GreB).

**Figure 3.**
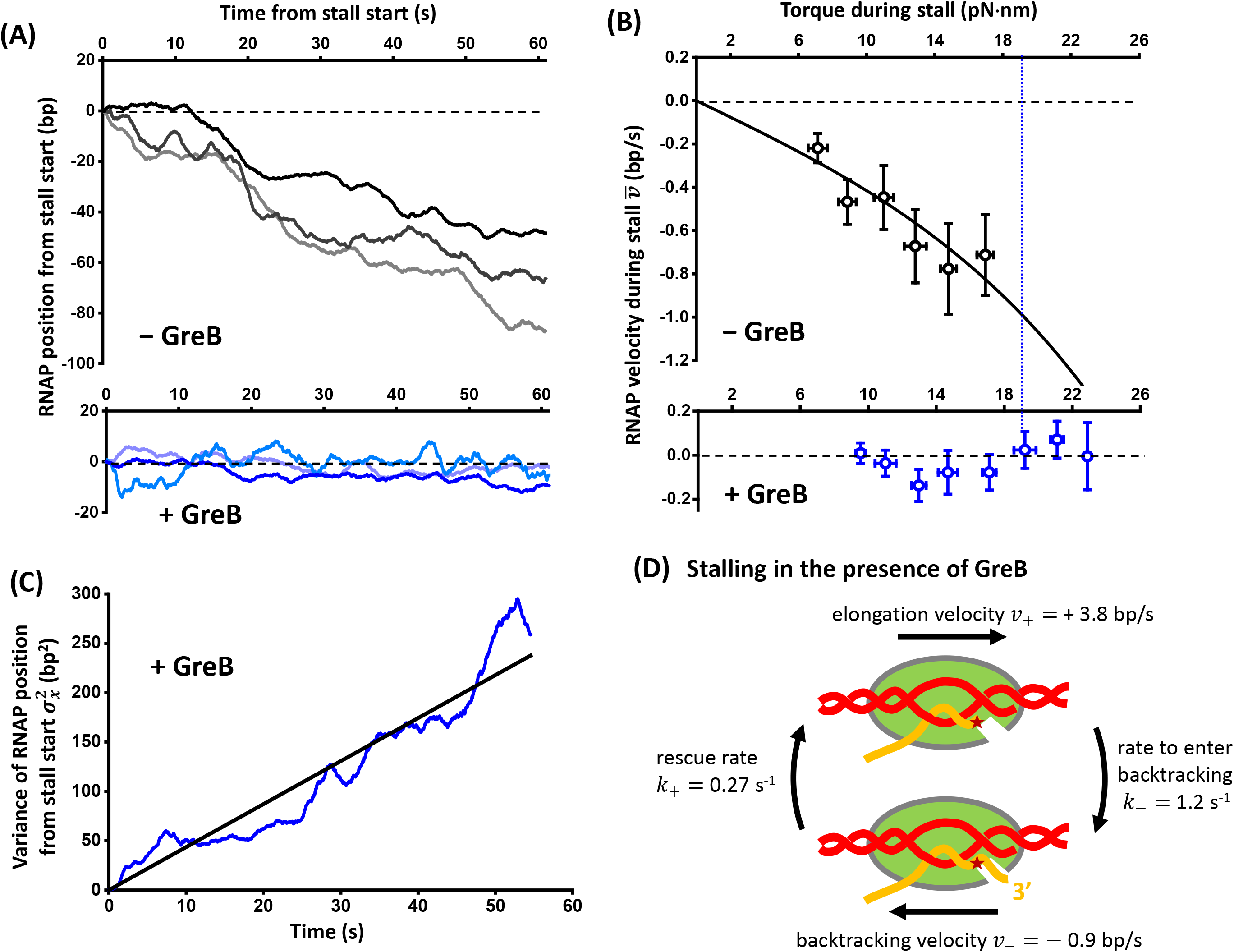
Kinetics of backtracking and GreB rescue. (**A**) Representative traces showing the RNAP position versus time after stalling in the absence (top) or presence of GreB (bottom). For clarity, all traces are aligned with respect to the start of stall in both time and position. These traces also have a torque ^∼^18 pN·nm upon stalling, close to the mean stall torque with GreB. (**B**) RNAP velocity versus torque during stall in the absence (top) and presence of GreB (bottom). The error bars of torque are SD and the error bars for RNAP velocity are SEMs. The black solid line is a fit to Eq. S1 (Supplementary Information), yielding a backtracking hopping rate in the absence of torque *k*_0_ = 0.26 s^-1^. The dashed line indicates the mean stall torque in the presence of GreB (18.5 pN·nm). Under this torque, the RNAP velocity in the absence of GreB represents the backtracking velocity *v_-_* = −0.9 bp/s. (**C**) Measured variance of RNAP position from the stall start versus time in the presence of GreB (blue). The black solid line is a linear fit passing through origin, yielding a slope of 4.4 bp^2^/s. (**D**) A two-state model of the RNAP during a stall in the presence of GreB (Supporting Information Text). Under a high resisting torque during a stall, an elongating RNAP may stochastically transition to a backtracking state. A backtracked RNAP may be rescued by GreB and transition back to an elongation state. We model the transitions between the two states as reversible first-order reactions. The lifetime of both states and the velocities of RNAP under both states determine the time course of RNAP’s position. See main text for the kinetic parameter values.

We also determined the active elongation velocity *v_+_* as a function of torque (Fig. S2; Methods). This was achieved by measuring the velocity between pauses prior to stalling. As expected, the active elongation velocity decreased with an increase in torque. In the absence of torque, RNAP translocated at around 20 bp/s under our condition (17, 40); whereas under a torque of 18.5 pN·nm, *v_+_* = +3.8 bp/s, both in the absence and presence of GreB (Fig. S2).

When RNAP stalled in the presence of GreB, RNAP’s motion fluctuated slightly around the initial stalling position (Fig. 3A, bottom), with the RNAP velocity 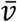 having minimal dependence on torque (Fig. 3B, bottom). The mean velocity was 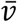 = 0.05 bp/s (Fig. S3), which has a speed much slower than that of backtracking or active elongation and has a magnitude comparable to the instrument’s drift speed (Fig. S1C).

A linear fit of the variance of RNAP position at a stall 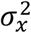 as a function of time yielded a slope of 4.4 bp^2^/s (Fig. 3C).

Taking all these together, the two-state model allowed us to obtain the GreB rescue rate to be *k_+_* = 0.27 s^-1^ and the rate to enter backtracking to be *k_-_* = 1.2 s^-1^ (Fig. 3D). Therefore, our analysis shows that upon stalling in the presence of GreB, RNAP backtracked ^∼^ 3 bp over ^∼^ 4 s before being rescued by GreB under the conditions used here. Our rescue rate is also in good agreement with that reported from a recent study of GreB which was performed under zero torsion and tension (45). This agreement suggests that GreB’s ability to rescue a backtracked RNAP may be insensitive to the torsion that RNAP experiences. Our theoretical model and the use of variance analysis thus provide a framework to obtain kinetic parameters from complex translocation trajectories.

### GreB Facilitates Transcription Resumption after a Stall

Upon torque relaxation, a stalled RNAP may exit the backtracked state to resume transcription, but how does the presence of GreB alter the rate of transcription resumption? To investigate, we first stalled the RNAP under torque, and then relaxed the torque to examine how fast transcription resumed, both in the absence and presence of GreB. As shown in Fig. 4, in the absence of GreB, ^∼^ 20% of stalled RNAPs resumed transcription within 35 s, consistent with our previous findings (17). In the presence of GreB, ^∼^ 40% resumed within 35 s. GreB also marginally increased the fraction of RNAPs that resumed transcription at longer time scales. Therefore, upon torque relaxation, GreB expedites transcription resumption. This is likely due to a combined effect of GreB shortening the backtracking distance during the initial stall and GreB rescuing backtracked complex upon torque relaxation.

**Figure 4.**
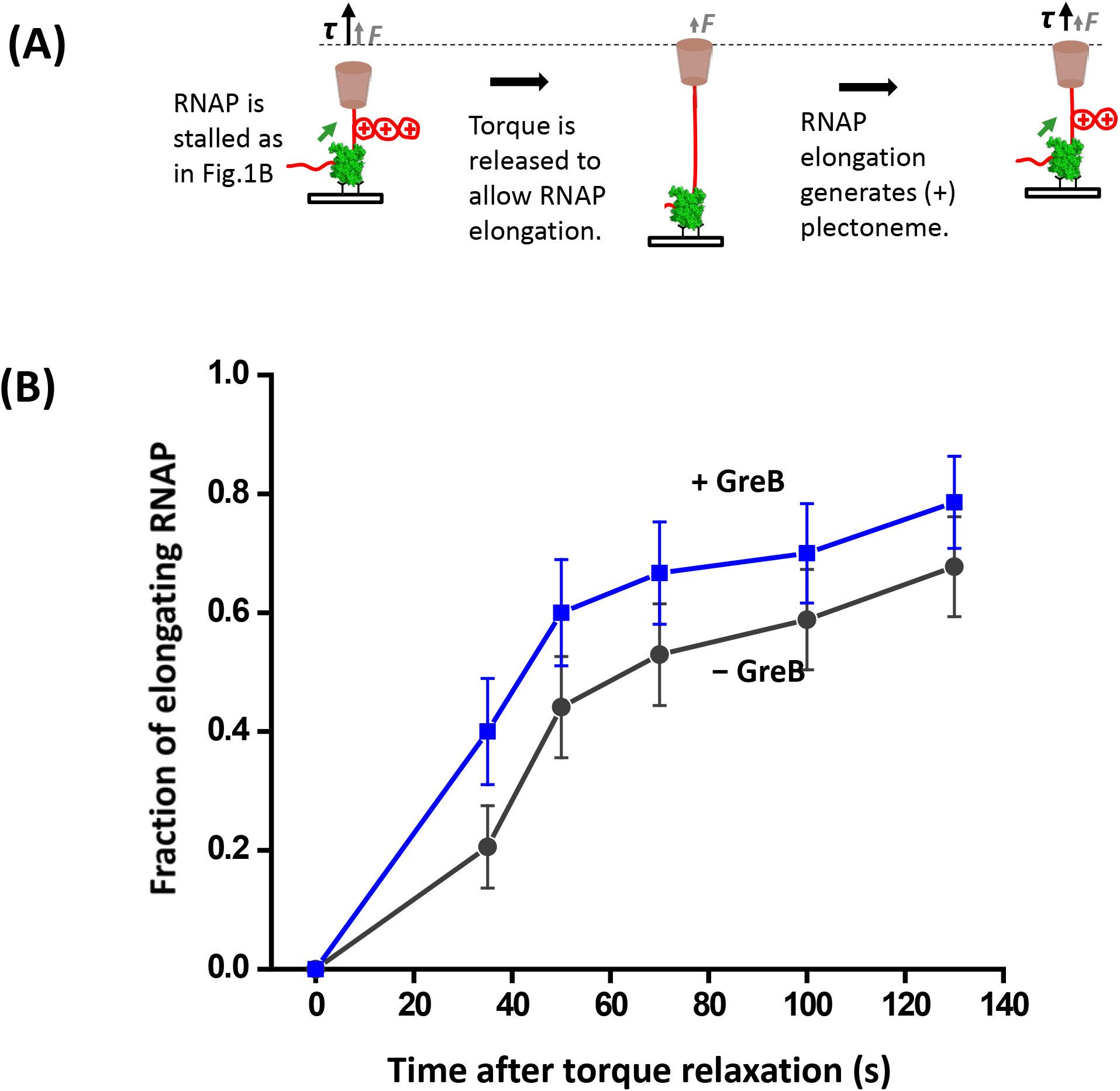
Transcription resumption in the presence of GreB. After RNAP was stalled, torque on the RNAP was relaxed to allow transcription resumption. Subsequent transcription was detected by repeating the RNAP stall torque measurement as shown in Fig. 1B. Error bars indicate standard error of means (SEM). Shown are the fractions of RNAPs that resumed transcription after torque release as a function of the time after torque relaxation.

**Figure 5.**
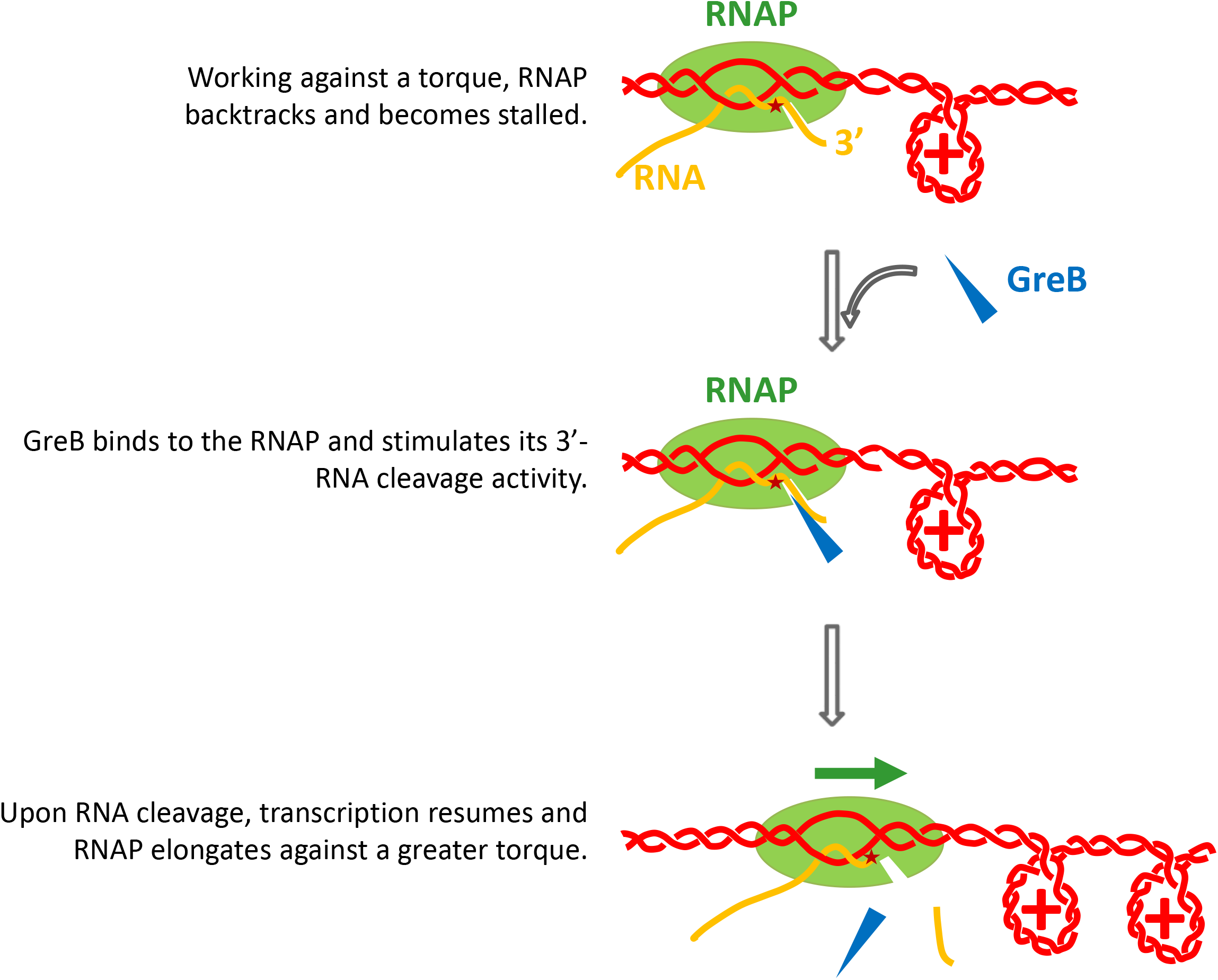
A proposed model for RNAP’s enhanced torsional capacity by GreB. During transcription, torsional stress accumulates with transcription and RNAP backtracks and is ultimately stalled. GreB can bind to the backtracked RNAP, stimulate its intrinsic RNA cleavage activity, and bring the RNAP back to elongation. Once transcription is resumed, the RNAP will be able to work against a stronger torsional barrier.

### Backtracking Arrest and Rescue” Model

Our experimental results support a “backtracking arrest and rescue” model for transcription under torsion in the presence of GreB (see Fig. 6). Torsional stress accumulated during transcription hinders RNAP forward translocation and is highly effective in inducing RNAP backtracking, which may lead to transcription arrest. GreB can bind to RNAP, interact with its secondary channel, stimulate RNAP’s cleavage activity, and thus prevent RNAP from being extensively backtracked. Consequently, the stalled RNAP can be rescued and transcription can be resumed. As this cycle repeats, RNAP works against an increasingly greater torque. RNAP ultimately stalls when GreB’s rescue of backtracking can no longer afford the RNAP with sufficient forward elongation to rectify backtracking.

## DISCUSSION

The work presented here demonstrates that backtracking is the primary mechanism that limits RNAP’s torsional capacity. This limitation can be mitigated by the transcription factor GreB which can decrease backtracking and enhance the torsional capacity of RNAP by 65%. *In vivo*, many other transcription factors may interact with RNAP to regulate RNAP’s torsional capacity as well. For example, NusA can assist RNAP backtracking (30) and, thus, potentially decrease RNAP’s stall torque, while factors like GreA (35), DksA (48), and NusG (29) prevent backtracking and may increase RNAP’s stall torque. These factors may work in coordination to downregulate or upregulate the stall torque of RNAP’s torsional capacity to achieve a broad range of regulation. Therefore, backtracking may serve as a general mediator for transcription factors to regulate RNAP torsional properties. In the cell, the enhanced torsional capacity may be necessary for overcoming obstacles such as removing road-block proteins (13) or transcribing a long gene under high torsional stress while ensuring continuous elongation. For example, *E. coil* RNAP must transcribe through various bound proteins such as HU and H-NS that must be displaced during transcription.

Although this work focuses on a prokaryotic RNAP, the findings here may have implications for its eukaryotic counterparts. Eukaryotic Pol II must transcribe through nucleosomes to access genetic information and nucleosomes form formidable barriers to transcription. It has recently been shown that transcription generated (+) torsional stress increases nucleosome turnover within genes (49) and indeed, single molecule studies found that a (+) torque of ^∼^19 pN·nm or larger can significantly disrupt nucleosome structures, leading to almost complete loss of H2A-H2B dimers (13). In addition, TFIIS, a functional analog of GreB in eukaryotic cells, has been shown to enhance the stall force of Pol II (2) and thus may also increase torsional capacity of Pol II and facilitate its passage through the nucleosome barrier.

This work has demonstrated that the RNAP’s torsional capacity can be regulated in a transcription factor-dependent manner. The interaction kinetics of GreB and RNAP are on the scale of base pairs and seconds. This type of transient interaction between proteins and factors must occur broadly in the cell but is difficult to capture due to the stochastic nature. The theoretical framework and experimental approach established here can potentially be applied to elucidate many other kinetic and mechanical processes that are important for cellular functions.

## METHODS

### DNA Preparations

Two DNA templates were prepared for the single-molecule experiments. A “downstream” DNA template was used for real-time single-molecule transcription assays to measure the downstream stall torque; while a “calibration” DNA template was used in the control experiment to measure the drift velocity. Both DNA templates contain a T7A1 promoter, and the preparation of these two templates was reported previously (17). For the “downstream” DNA template, a 4.7-kb DNA segment was amplified by PCR from the plasmid pMDW2 (sequence available upon request) and digested via BasI-HF at 37 °C for 1 hour to generate a 4-bp 5’-overhang downstream of the T7A1 promoter. The digested fragment was then ligated to a 66-bp DNA torsional-constraint anchor with 6 biotin linkers evenly spaced for attachment to the streptavidin-coated bottom surface of a quartz cylinder at 16 °C overnight. The “calibration” template was the same as the “downstream” template except that the upstream end of the T7A1 promoter was also digested to generate a 4-bp 5’-overhang and ligated to a 6-digoxigenin linker with dig tags evenly spaced for attachment to an anti-dig coated coverslip surface. The sequences of these two templates are available upon request.

### Purification of *E. coli* RNAP

*E. coli* RNAP containing one hemagglutinin (HA) epitope tag on the C-terminus of each α-subunit was expressed and purified as previously described (50, 51). Briefly, RNAP was expressed at low levels in DH-5α-competent *E. coli* transformed with the plasmid pKA1 in Superbroth with 100 μg/ml ampicillin at 37 °C until OD_600nm_ reached 2.1. IPTG was added to a final concentration of 1 mM, and the cells were grown for an additional 4 h, harvested by centrifugation and stored at −20 °C. The transformed cells were re-suspended at 3 ml/g in RNAP Lysis Buffer (50 mM Tris-HCl pH 8.0, 300 mM NaCl, 10 mM EDTA, 1 mM DTT, and 5% v/v glycerol). Lysozyme was added to 300 μg/ml, and the cells were incubated for 20 min on ice. Further lysis was achieved by sonication via a Branson Sonifier 250 with 60% duty cycle in small aliquots (< 20 ml). The cell debris was pelleted by centrifugation at 16,300 g for 45 min at 4 °C and the supernatant containing DNA and DNA-bound proteins was collected. Cleared 5% (w/v) polyethyleneimine (PEI) was added dropwise, with slow stirring, to a final concentration of 0.4% (w/v) in order to precipitate nucleic acids and their bound proteins. The DNA and associated RNAP were pelleted from solution and after five washes in a buffer containing 350 mM NaCl, RNAP was eluted from PEI and DNA with a buffer containing 1 M NaCl. The eluted RNAP was purified to homogeneity through chromatography on 3 columns: first on a HiPrep Heparin FF 16/10 column (GE), followed by a HiPrep 26/60 Sephacryl S-300 HR column (GE), and last on a Ni-NTA Superflow column (QIAGEN). Fractions containing pure RNAP were pooled, concentrated and dialyzed against RNAP Storage Buffer (50 mM Tris-HCl pH 8.0, 100 mM NaCl, 1 mM EDTA, 1 mM DTT, and 50% v/v glycerol) and stored at −20 °C.

### Purification of GreB

Plasmid pES3, encoding GreB-6xHis in pET-28b(+) (37), was transformed into BL21(DE3) cells for protein overexpression. Cells were grown in LB media supplemented with 50 μg/ml of kanamycin at 37 °C until the OD_600nm_ reached 0.6-0.8, and then were induced with 1 mM IPTG. After induction at 37 °C for 3 hours, cells were harvested and stored at −80 °C. The protocol for the purification of GreB proteins was similar to those previous described (37, 52). In brief, the cells were thawed on ice and re-suspended in GreB Lysis Buffer (50 mM Tris-HCl pH 6.9, 500 mM NaCl, and 5% v/v glycerol) with EDTA-free protease-inhibitor cocktail (Roche) and lysozyme (300 μg/ml). The suspension was incubated on ice for 1 hour and followed by a brief sonication to further lyse the cells. The extract was cleared by centrifugation (24,000 g, 20 min at 4 °C) and passed twice through a 0.45-μm filter. The proteins were isolated by a Ni-NTA agarose column and eluted in GreB Lysis Buffer with 200 mM imidazole. The elute was further purified on a Superdex 200 column (GE Healthcare) with Elution Buffer (10 mM Tris-HCl pH 8.0, 500 mM NaCl, 1 mM DTT, 1 mM EDTA, and 5% v/v glycerol). The purified proteins were dialyzed into GreB storage buffer (10 mM Tris-HCl pH 8.0, 200 mM NaCl, 1 mM DTT, 1 mM EDTA, and 50% v/v glycerol), frozen in liquid nitrogen and stored at −80 °C.

### Paused Transcription Complexes Assembly

The transcription elongation complexes were formed in bulk and paused at +20 position via nucleotide depletion (17, 51). In brief, 25 nM HA-tagged *E. coli* RNAP was incubated with 5 nM “Downstream” DNA templates, 250 μM ApU (initiating dinucleotide), and 50 μM GTP/ATP/CTP in the Transcription Buffer (25 mM Tris-HCl pH 8.0, 100 mM KCl, 4 mM MgCfe, 1 mM DTT, and 3% v/v glycerol) with 1 unit/μl SUPERase-In^TM^ RNase inhibitor and 0.15 mg/ml acetylated BSA at 37 °C for 40 minutes. Heparin was then added to 0.2 μg/μl and the paused transcription complexes (PTCs) were kept at 4 ºC before being used in the single molecule assays.

### DNA supercoiling characterization

We performed a similar DNA supercoiling property characterization as described before in order to extract desired information (such as stall torque, RNAP template position, transcription velocity, etc.) from measured data (such as force, DNA extension, etc.) (17). These were conducted using the “calibration” DNA template in the transcription buffer (Fig. S4).

### Instrument drift estimate

A control experiment that mimics the real-time single-molecule transcription assay was employed to estimate instrument drift (Fig. S1). In brief, the “calibration” DNA had one end tonsionally anchored on a coverslip surface coated with anti-dig while the other end was torsionally constrained to the bottom surface of a quartz cylinder via multiple biotin/streptavidin linkages. The DNA was first stretched and mechanically wound by 12 turns under the force clamp of 0.3 pN to generate the (+) plectoneme. Then the force clamp was turned off. The trap position was held constant for 60 s to mimic the stall torque measurement step in the real-time transcription assay and the extension change was recorded. The drift velocity was obtained by the linear-fitting of the extension change.

### Real-Time Single-Molecule Transcription Assay

The angular-optical-trap-based single-molecule transcription assays were performed as previously described (17). Briefly, an RNAP in the PTC was torsionally anchored on a coverslip surface coated with anti-HA through two HA/anti-HA interactions; while the downstream end of DNA was torsionally constrained to the bottom surface of a quartz cylinder via multiple biotin/streptavidin linkages (Fig. 1A). After the introduction of 1 mM NTPs to resume transcription, DNA was mechanically unwound to form a (-) plectoneme under the force clamp of 0.3 pN (Fig. 1B). Subsequent translocation of RNAP neutralized the (-) plectoneme and led to the (+) plectoneme formation. The force clamp was then turned off. Subsequent RNAP translocation increased the resisting force and the corresponding torque until reaching a stall. For experiments in the presence of GreB factor, 1 μM of GreB was mixed with 1 mM NTPs to restart transcription elongation. These single-molecule transcription assays were carried out in an environmentally controlled room at 23.2 ± 0.5 °C.

### Data Acquisition and Analysis

During a transcription experiment, RNAP buckled the DNA and was allowed to stall under an increasing torque (Fig. 1A). The DNA contour length and torque as a function of time were obtained from the measured force, DNA extension, and cylinder orientation as a function of time as previously derived (17). Raw data for measured force (F), DNA extension (z), and cylinder orientation were low-pass filtered to 1 kHz and digitized at 2 kHz. The RNAP template position relative to the transcription start site was thus determined from the DNA contour length and the known transcription start site location. The stall torque was taken as the torque value averaged over 2 s upon stalling.

For pause-free velocity analysis, the converted RNAP template position data were smoothed using a Gaussian weight function with a standard deviation of 0.1 s and pauses were identified as previously reported (17). The cutoff duration was set to be 0.2 s (i.e., an RNAP was considered to have paused if the time it spent at a single nucleotide position was greater than 0.2 s). The 0.2 s was found to be 4 times the most likely dwell time which represented the baseline. Pause-free velocity was determined for the data segments between pauses.

To obtain RNAP velocity at a stall versus torque (Figure 3B), we took into account the decrease in torque as RNAP backtracked in each trace (see an example in Figure 1C). Each trace was first divided into 20 s segments, and for each segment, the RNAP velocity and the mean torque were determined. RNAP velocities from different segments of all traces were then binned according to their torque values before being averaged

## Acknowledgements

We thank the members of the Wang Laboratory for critical discussion and comments on this work. We especially thank Dr. James T. Inman for technical assistance with the angular optical trap, Dr. Eric Strobel for sharing the GreB plasmid and purification protocol, and Drs. James P. Sethna and Anatoly Kolomeisky for helpful discussions on biased random walks. This work was supported by the Howard Hughes Medical Institute (to M.D.W.), the National Science Foundation grants (MCB-0820293 and MCB-1517764 to M.D.W.), the National Natural Science Foundation of China (NSFC-11674403 to J.M.), and the Fundamental Research Funds for the Central Universities (15lgjc15 to J.M.).

## References

1. Wang MD, et al. (1998) Force and velocity measured for single molecules of RNA polymerase. Science 282(5390):902–907.

2. Galburt EA, et al. (2007) Backtracking determines the force sensitivity of RNAP II in a factor-dependent manner. Nature 446(7137):820–823.

3. Ma J & Wang MD (2014) RNA polymerase is a powerful torsional motor. Cell Cycle 13(3):337–338.

4. Liu LF & Wang JC (1987) Supercoiling of the DNA template during transcription. Proc Natl Acad Sci U S A 84(20):7024–7027.

5. Kouzine F, et al. (2013) Transcription-dependent dynamic supercoiling is a short-range genomic force. Nature Structural & Molecular Biology 20(3):396–403.

6. Naughton C, et al. (2013) Transcription forms and remodels supercoiling domains unfolding large-scale chromatin structures. Nat Struct Mol Biol 20(3):387–395.

7. Teves SS & Henikoff S (2014) Transcription-generated torsional stress destabilizes nucleosomes. Nature Structural & Molecular Biology 21(1):88–94.

8. Leng FF, Amado L, & McMacken R (2004) Coupling DNA supercoiling to transcription in defined protein systems. Journal of Biological Chemistry 279(46):47564–47571.

9. Oberstrass FC, Fernandes LE, & Bryant Z (2012) Torque measurements reveal sequence-specific cooperative transitions in supercoiled DNA. Proc Natl Acad Sci U S A 109(16):6106–6111.

10. Ding Y, et al. (2014) DNA supercoiling: A regulatory signal for the lambda repressor. Proceedings of the National Academy of Sciences of the United States of America 111(43):15402–15407.

11. Kouzine F, Liu JH, Sanford S, Chung HJ, & Levens D (2004) The dynamic response of upstream DNA to transcription-generated torsional stress. Nature Structural & Molecular Biology 11(11):1092–1100.

12. Kouzine F, Sanford S, Elisha-Feil Z, & Levens D (2008) The functional response of upstream DNA to dynamic supercoiling in vivo. Nature Structural & Molecular Biology 15(2):146–154.

13. Sheinin MY, Li M, Soltani M, Luger K, & Wang MD (2013) Torque modulates nucleosome stability and facilitates H2A/H2B dimer loss. Nature Communications 4.

14. Harada Y, et al. (2001) Direct observation of DNA rotation during transcription by Escherichia coli RNA polymerase. Nature 409(6816):113–115.

15. Matsumoto K & Hirose S (2004) Visualization of unconstrained negative supercoils of DNA on polytene chromosomes of Drosophila. J Cell Sci 117(Pt 17):3797–3805.

16. Wu HY, Shyy SH, Wang JC, & Liu LF (1988) Transcription generates positively and negatively supercoiled domains in the template. Cell 53(3):433–440.

17. Ma J, Bai L, & Wang MD (2013) Transcription Under Torsion. Science 340(6140):1580–1583.

18. Lipfert J, Kerssemakers JWJ, Jager T, & Dekker NH (2010) Magnetic torque tweezers: measuring torsional stiffness in DNA and RecA-DNA filaments. Nature Methods 7(12):977–U954.

19. Oberstrass FC, Fernandes LE, Lebel P, & Bryant Z (2013) Torque Spectroscopy of DNA: Base-Pair Stability, Boundary Effects, Backbending, and Breathing Dynamics. Physical Review Letters 110(17).

20. Sheinin MY, Forth S, Marko JF, & Wang MD (2011) Underwound DNA under Tension: Structure, Elasticity, and Sequence-Dependent Behaviors. Physical Review Letters 107(10).

21. Lilley DMJ & Higgins CF (1991) Local DNA topology and gene expression: the case of the leu-500 promoter. Molecular Microbiology 5(4):779–783.

22. Revyakin A, Ebright RH, & Strick TR (2004) Promoter unwinding and promoter clearance by RNA polymerase: detection by single-molecule DNA nanomanipulation. Proc Natl Acad Sci U S A 101(14):4776–4780.

23. Revyakin A, Liu C, Ebright RH, & Strick TR (2006) Abortive initiation and productive initiation by RNA polymerase involve DNA scrunching. Science 314(5802):1139–1143.

24. Bryant Z, et al. (2003) Structural transitions and elasticity from torque measurements on DNA. Nature 424(6946):338–341.

25. Deufel C, Forth S, Simmons CR, Dejgosha S, & Wang MD (2007) Nanofabricated quartz cylinders for angular trapping: DNA supercoiling torque detection. Nat Methods 4(3):223–225.

26. Forth S, et al. (2008) Abrupt buckling transition observed during the plectoneme formation of individual DNA molecules. Physical Review Letters 100(14).

27. Forth S, Sheinin MY, Inman J, & Wang MD (2013) Torque measurement at the single-molecule level. Annu Rev Biophys 42:583–604.

28. Borukhov S, Lee J, & Laptenko O (2005) Bacterial transcription elongation factors: new insights into molecular mechanism of action. Molecular Microbiology 55(5):1315–1324.

29. Herbert KM, et al. (2010) E. coli NusG Inhibits Backtracking and Accelerates Pause-Free Transcription by Promoting Forward Translocation of RNA Polymerase. Journal of Molecular Biology 399(1):17–30.

30. Zhou J, Ha Kook S, La Porta A, Landick R, & Block Steven M (2011) Applied Force Provides Insight into Transcriptional Pausing and Its Modulation by Transcription Factor NusA. Molecular Cell 44(4):635–646.

31. Komissarova N & Kashlev M (1997) Transcriptional arrest: Escherichia coli RNA polymerase translocates backward, leaving the 3’ end of the RNA intact and extruded. Proc Natl Acad Sci U S A 94(5):1755–1760.

32. Nudler E, Mustaev A, Lukhtanov E, & Goldfarb A (1997) The RNA-DNA hybrid maintains the register of transcription by preventing backtracking of RNA polymerase. Cell 89(1):33–41.

33. Izban MG & Luse DS (1993) SII-facilitated transcript cleavage in RNA polymerase II complexes stalled early after initiation occurs in primarily dinucleotide increments. J Biol Chem 268(17):12864–12873.

34. Reinberg D & Roeder RG (1987) Factors involved in specific transcription by mammalian RNA polymerase II. Purification and functional analysis of initiation factors IIB and IIE. J Biol Chem 262(7):3310–3321.

35. Marr MT & Roberts JW (2000) Function of transcription cleavage factors GreA and GreB at a regulatory pause site. Molecular Cell 6(6):1275–1285.

36. Stepanova E, Wang M, Severinov K, & Borukhov S (2009) Early transcriptional arrest at Escherichia coli rplN and ompX promoters. J Biol Chem 284(51):35702–35713.

37. Strobel EJ & Roberts JW (2014) Regulation of promoter-proximal transcription elongation: enhanced DNA scrunching drives lambda Q antiterminator-dependent escape from a sigma 70-dependent pause. Nucleic Acids Research 42(8):5097–5108.

38. Chedin S, Riva M, Schultz P, Sentenac A, & Carles C (1998) The RNA cleavage activity of RNA polymerase III is mediated by an essential TFIIS-like subunit and is important for transcription termination. Genes Dev 12(24):3857–3871.

39. La Porta A & Wang MD (2004) Optical torque wrench: Angular trapping, rotation, and torque detection of quartz microparticles. Physical Review Letters 92(19).

40. Bai L, Fulbright RM, & Wang MD (2007) Mechanochemical kinetics of transcription elongation. Phys Rev Lett 98(6):068103.

41. Yin H, et al. (1995) Transcription against an applied force. Science 270(5242):1653–1657.

42. Shaevitz JW, Abbondanzieri EA, Landick R, & Block SM (2003) Backtracking by single RNA polymerase molecules observed at near-base-pair resolution. Nature 426(6967):684–687.

43. Herbert KM, et al. (2006) Sequence-resolved detection of pausing by single RNA polymerase molecules. Cell 125(6):1083–1094.

44. Shundrovsky A, Santangelo TJ, Roberts JW, & Wang MD (2004) A single-molecule technique to study sequence-dependent transcription pausing. Biophys J 87(6):3945–3953.

45. Tetone LE, et al. (2017) Dynamics of GreB-RNA polymerase interaction allow a proofreading accessory protein to patrol for transcription complexes needing rescue. Proc Natl Acad Sci U S A 114(7): E1081–E1090.

46. Bai L, Shundrovsky A, & Wang MD (2004) Sequence-dependent kinetic model for transcription elongation by RNA polymerase. J Mol Biol 344(2):335–349.

47. Depken M, Galburt EA, & Grill SW (2009) The origin of short transcriptional pauses. Biophys J 96(6):2189–2193.

48. Zhang Y, et al. (2014) DksA Guards Elongating RNA Polymerase against Ribosome-Stalling-Induced Arrest. Molecular Cell 53(5):766–778.

49. Teves SS & Henikoff S (2014) Transcription-generated torsional stress destabilizes nucleosomes. Nat Struct Mol Biol 21(1):88–94.

50. Le TT, et al. (2018) Mfd Dynamically Regulates Transcription via a Release and Catch-Up Mechanism. Cell 172(1-2):344–357 e315.

51. Adelman K, et al. (2002) Single molecule analysis of RNA polymerase elongation reveals uniform kinetic behavior. Proceedings of the National Academy of Sciences of the United States of America 99(21):13538–13543.

52. Perederina AA, et al. (2006) Cloning, expression, purification, crystallization and initial crystallographic analysis of transcription elongation factors GreB from Escherichia coli and Gfh1 from Thermus thermophilus. Acta Crystallographica Section F-Structural Biology and Crystallization Communications 62:44–46.

